# scPipe: a flexible R/Bioconductor preprocessing pipeline for single-cell RNA-sequencing data

**DOI:** 10.1101/175927

**Authors:** Luyi Tian, Shian Su, Xueyi Dong, Daniela Amann-Zalcenstein, Christine Biben, Azadeh Seidi, Douglas J. Hilton, Shalin H. Naik, Matthew E. Ritchie

**Affiliations:** Molecular Medicine Division, The Walter and Eliza Hall Institute of Medical Research, 1G Royal Parade, Parkville, 3052, Australia; Department of Medical Biology, The University of Melbourne, Parkville, 3010, Australia; College of Life Science, Zhejiang University, 866 Yuhangtang Road, Hangzhou, Zhejiang Province, 310058, P.R. China; Australian Genome Research Facility, 1G Royal Parade, Parkville, 3052, Australia; School of Mathematics and Statistics, The University of Melbourne, Parkville, 3010, Australia

## Abstract

Single-cell RNA sequencing (scRNA-seq) technology allows researchers to profile the transcriptomes of thousands of cells simultaneously. Protocols that incorporate both designed and random barcodes have greatly increased the throughput of scRNA-seq, but give rise to a more complex data structure. There is a need for new tools that can handle the various barcoding strategies used by different protocols and exploit this information for quality assessment at the sample-level and provide effective visualization of these results in preparation for higher-level analyses.

To this end, we developed *scPipe*, a R/Bioconductor package that integrates barcode demultiplexing, read alignment, UMI-aware gene-level quantification and quality control of raw sequencing data generated by multiple 3-prime-end sequencing protocols that include CEL-seq, MARS-seq, Chromium 10X and Drop-seq. *scPipe* produces a count matrix that is essential for downstream analysis along with an HTML report that summarises data quality. These results can be used as input for downstream analyses including normalization, visualization and statistical testing. *scPipe* performs this processing in a few simple R commands, promoting reproducible analysis of single-cell data that is compatible with the emerging suite of scRNA-seq analysis tools available in R/Bioconductor. The *scPipe* R package is available for download from https://www.bioconductor.org/packages/scPipe.

## Introduction

Advances in single-cell transcriptomic profiling technologies allow researchers to measure gene activity in thousands of cells simultaneously, enabling exploration of gene expression variability [1], identification of new cell types [2] and the study of transcriptional programs involved in cell differentiation [3]. The introduction of cellular barcodes, sequences distinct for each cell attached to the dT-primer, has increased the throughput and substantially reduced the cost of single-cell RNA sequencing (scRNA-seq). These barcodes allow for the demultiplexing of reads after cells are pooled together for sequencing. Apart from cellular barcodes, molecular barcodes or unique molecular identifiers (UMIs), are frequently employed to remove PCR duplicates and allow identification of unique mRNA molecules, thereby reducing technical noise. The multiple levels of barcoding used in scRNA-seq experiments create additional challenges in data processing together with new opportunities for quality control (QC). Different protocols use different barcode configurations, which means a flexible approach to data preprocessing is required.

A large number of analysis tools have already been tailored to scRNA-seq analysis [4], the majority of which are focused on downstream analysis tasks such as clustering and trajectory analysis. Methods for preprocessing tend to focus on specific tasks such as *UMI-tools* [5], *umitools* (http://brwnj.github.io/umitools/) *and umis* [6] which have been developed for handling random UMIs and correcting UMI sequencing errors. Other tools such as *CellRanger* [7], *dropEst* [8] and *dropseqPipe* (https://github.com/Hoohm/dropSeqPipe) on the other hand offer a complete preprocessing solution for data generated by droplet based protocols. Other packages such as *scater* [9], *and scran* [10] work further downstream by preprocessing the counts to perform general QC and normalization of scRNA-seq data.

*scPipe* was developed to address the lack of a comprehensive R-based workflow for processing sequencing data from 3-prime (3‘) end protocols that can accommodate both UMIs and sample barcodes, map reads to the genome and summarise these results into gene-level counts. Additionally this pipeline collates useful metrics for QC during preprocessing that can be later used to filter genes and samples. In the remainder of this article we provide an overview of the main features of our *scPipe* software and demonstrate its use on two scRNA-seq datasets.

## Materials and Methods

### Single cell RNA-seq datasets

#### Mouse hematopoietic lineage dataset

Single cell expression profiling of the main hematopoietic lineages in mouse (erythroid, myeloid, lymphoid, stem/progenitors) was performed using a modified CEL-seq2 [11] protocol. B lymphocytes (B220+ FSC-Alow), erythroblasts (Ter119+ CD44+, FSC-Amid/high), granulocytes (Mac1+ Gr1+) and high-end progenitor/stem (Lin-Kit+ Sca1+) were sorted from the bone marrow of a C57BL/6 10-13 week old female mouse. T cells (CD3+ FSC-Alow) were isolated from the thymus of the same mouse. Bone marrow and thymus were dissociated mechanically, washed and stained with antibodies for 1hr on ice. Single cells were deposited on a 384 well plate using an Aria cell sorter (Beckman). Index data was collected and our adapted CEL-seq2 protocol used to generate a library for sequencing. The reads were sequenced by an Illumina Nextseq 500 and processed by *scPipe*. This dataset is available under GEO accession number GSE109999.

#### Human lung adenocarcinoma cell line dataset

The cell lines H2228, NCI-H1975 and HCC827 were retrieved from ATCC (https://www.atcc.org/) and cultured in Roswell Park Memorial Institute (RPMI) 1640 medium with 10% fetal calf serum (FCS) and 1% Penicillin-Streptomycin. The cells were grown independently at 37°C with 5% carbon dioxide until near 100% confluency. Cells were PI stained and 120,000 live cells were sorted for each cell line by FACS to acquire an accurate equal mixture of live cells from the three cell lines. This mixture was then processed by Chromium 10X single cell platform using the manufacturer’s (10X Genomics) protocol and sequenced with an Illumina Nextseq 500. Filtered gene expression matrices were generated using *CellRanger* (10X Genomics) and *scPipe* independently. For *CellRanger*, we used the default parameters, with `--expect-cells=4000`. For *scPipe*, we processed the 4000 most enriched cell barcodes. The GRCh38 human genome and associated annotation were used for both the *scPipe* and *CellRanger* analysis. This dataset is available under GEO accession number GSE111108.

### Results and Discussion

#### Implementation

*scPipe* is an R [12] / Bioconductor [13] package that can handle data generated from all popular 3’ end scRNA-seq protocols and their variants, such as CEL-seq, MARS-seq, Chromium 10X and Drop-seq. Although not specifically designed for non-UMI protocols, it can also process data generated by Smart-seq and Smart-seq2. The pipeline begins with FASTQ files and outputs both a gene count matrix and a variety of QC statistics. These are presented in a standalone HTML report generated by *rmarkdown* [14] *that includes various plots of QC metrics and other data summaries. The scPipe* package is written in R and C++ and uses the *Rcpp* package [15, 16] to wrap the C++ code into R functions and the *Rhtslib* package [17] for BAM input/output. The key aspects are implemented in C++ for efficiency. The *scPipe* package is available from https://www.bioconductor.org/packages/scPipe.

### The *scPipe* workflow

#### FASTQ reformatting

The *scPipe* workflow (Figure 1) begins with paired-end FASTQ data which is passed to the function `sc_trim_barcode`, which reformats the reads by trimming the barcode and UMI from the reads and moving this information into the read header. There are options to perform some basic filtering in this step, including removing reads with low quality sequence in the barcode and UMI regions and filtering out low complexity reads, most of which are non-informative repetitive sequences such as polyA. The output FASTQ file contains transcript sequences, with barcode information merged into the read names.

**Fig 1.**
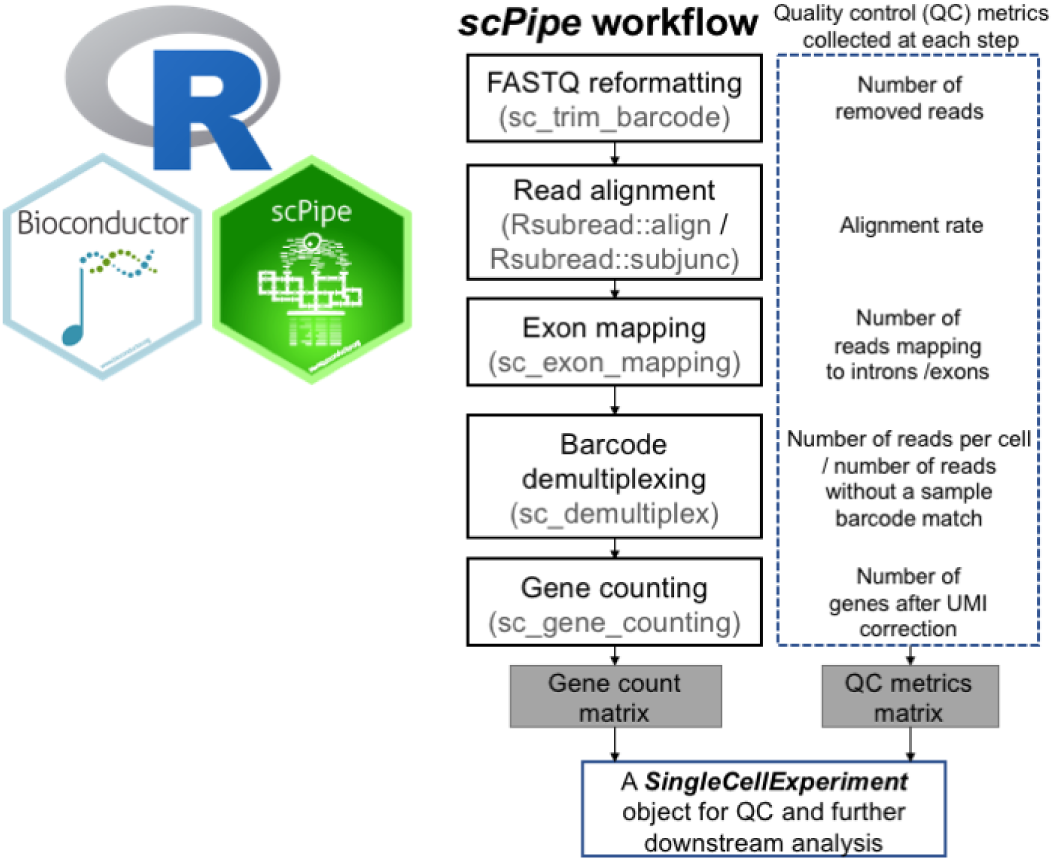
***The scPipe* workflow.** *scPipe* is a R/Bioconductor package that uses functionality from a number of other packages, including *Rsubread* to align reads to a reference genome (although in practice any aligner that produces BAM files can be used for read alignment) and *SingleCellExperiment* to organise the counts and sample annotation information. The major steps in the preprocessing pipeline of *scPipe* are shown along with the quality control (QC) statistics collected at each stage. The final output of this process is a matrix of counts and QC metrics for use in downstream analysis.

#### Read alignment, exon mapping and barcode demultiplexing

The next stage of preprocessing involves sequence alignment. For this task, any popular RNA-seq read aligner that produces BAM formatted output can be used. By default, *scPipe* uses the fast and convenient R-based *Rsubread* [18] aligner which is available on Linux and Mac OS operating systems. Aligned reads in the BAM file are then assigned to the exons by the sc exon mapping function according to a user provided annotation. Using a similar strategy to featureCounts [19], we divide chromosomes into non-overlapping bins and assess the overlap of aligned fragments with exons that fall within the bin to reduce the search complexity. This function records the mapping result, together with the UMI and cell barcodes available from the optional fields of the BAM file with specific BAM tags. By default we use the official BAM tag BC for cell barcode and OX for UMI sequence. Next, the `sc_demultiplex` function is used to demultiplex results per cell using the sample barcode information. For CEL-seq and MARS-seq, the demultiplexing is based on the list of designed barcode sequences that the user provides, and allows for mismatches during the cell barcode matching step. For 10X and Drop-seq where the cell barcode sequences need to be identified from the reads, the function `sc_detect_bc` can be applied to identify enriched barcodes in the data that can then be supplied to `sc_demultiplex`. Data from non-UMI protocols such as Smart-seq and Smart-seq2 can be handled by setting `has_UMI` to `TRUE` when running `sc_demultiplex`. The `sc_detect_bc` function summarises the sequences in the cell barcode region, collapsing reads with a small edit distance, which are indicative of sequencing errors. It can report a given number of cell barcodes or keep the cell barcodes based on a read number cutoff. A relaxed threshold is generally recommended, for instance if the estimated number of cells in the library is 4,000, a threshold of 6,000 would ensure data for all cells is collected. The quality control function in *scPipe* can be used to remove erroneous cell barcodes at a later stage. The demultiplexed data are output in a csv file for each cell, where each row corresponds to a read that maps to a specific gene, and columns that include gene id, UMI sequence and mapping position. The overall barcode demultiplexing results are recorded and can be plotted as shown in Figure 2A for the mouse blood cells generated by the CEL-seq2 protocol.

**Fig 2.**
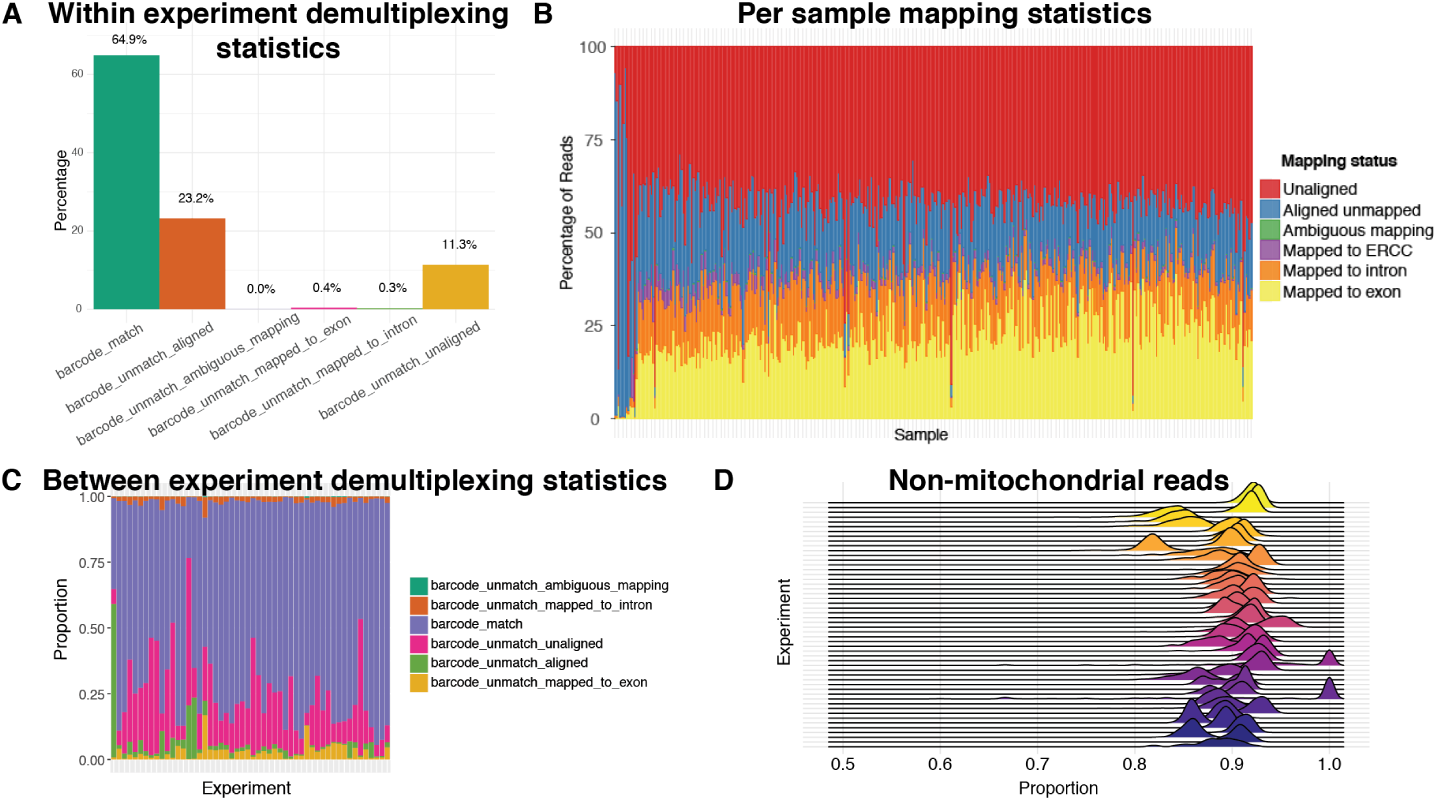
**Example QC plots that can be created using output from *scPipe* to assess data quality both within and between experiments.** Within experiment displays include (**A**) a bar plot illustrating the overall cell barcode matching results to assess sequencing accuracy across all samplaes and (**B**) a stacked bar plot showing the mapping rate, separated into reads that map to exon, intron and ERCCs and those that are ambiguously mapped, map elsewhere in the genome and are unaligned for each cell in an experiment (ordered by exon mapping rate). Between experiment displays include (**C**) a stacked bar plot showing the cell barcode matching results from panel (A) from multiple experiments and (**D**) a ridgeline plot presenting the distribution of proportions of non-mitochondrial read counts for cells across multiple experiments.

#### Obtaining a gene count matrix and performing quality assessment

The next stage of preprocessing makes use of the `sc_gene_counting` function to remove PCR duplicates (also known as UMI deduplication) to generate a gene count matrix. *scPipe* employs a simple method to correct for UMI sequencing errors by collapsing UMI sequences assigned to the same gene that have a low edit distance. After UMI deduplication we get a gene count matrix that can be used for further analysis. QC information is collected at each step (Figure 1) and includes the total number of mapped reads per cell, UMI deduplication statistics, per cell barcode demultiplexing statistics, UMI correction statistics and External RNA Controls Consortium (ERCC) spike-in control statistics (where present).

*scPipe* uses infrastructure from the *SingleCellExperiment* package [20] for data storage. A `SingleCellExperiment` object can be constructed by the `create_sce_by_dir` function using the data folder generated during preprocessing, or by manually specifying all the required slots. Alongside the gene count matrix, this object stores QC information obtained during preprocessing, and the type of gene id and organism name, which can both be useful in downstream functional enrichment analysis.

Next, alignment statistics for each cell can be plotted using the `plotMapping` function (Figure 2B). Data shown here is again from the mouse blood cell dataset described previously. Monitoring these QC statistics experiment-wide over time (Figures 2C-D) can be particularly useful for labs that routinely process single-cell data, allowing them to assess the impact changes in lab processes and protocols have on data quality. Pairwise scatter plots of QC metrics such as the total molecules per cell and the number of genes detected can be generated with the `plotQC_pair` function (Figure 3).

**Fig 3.**
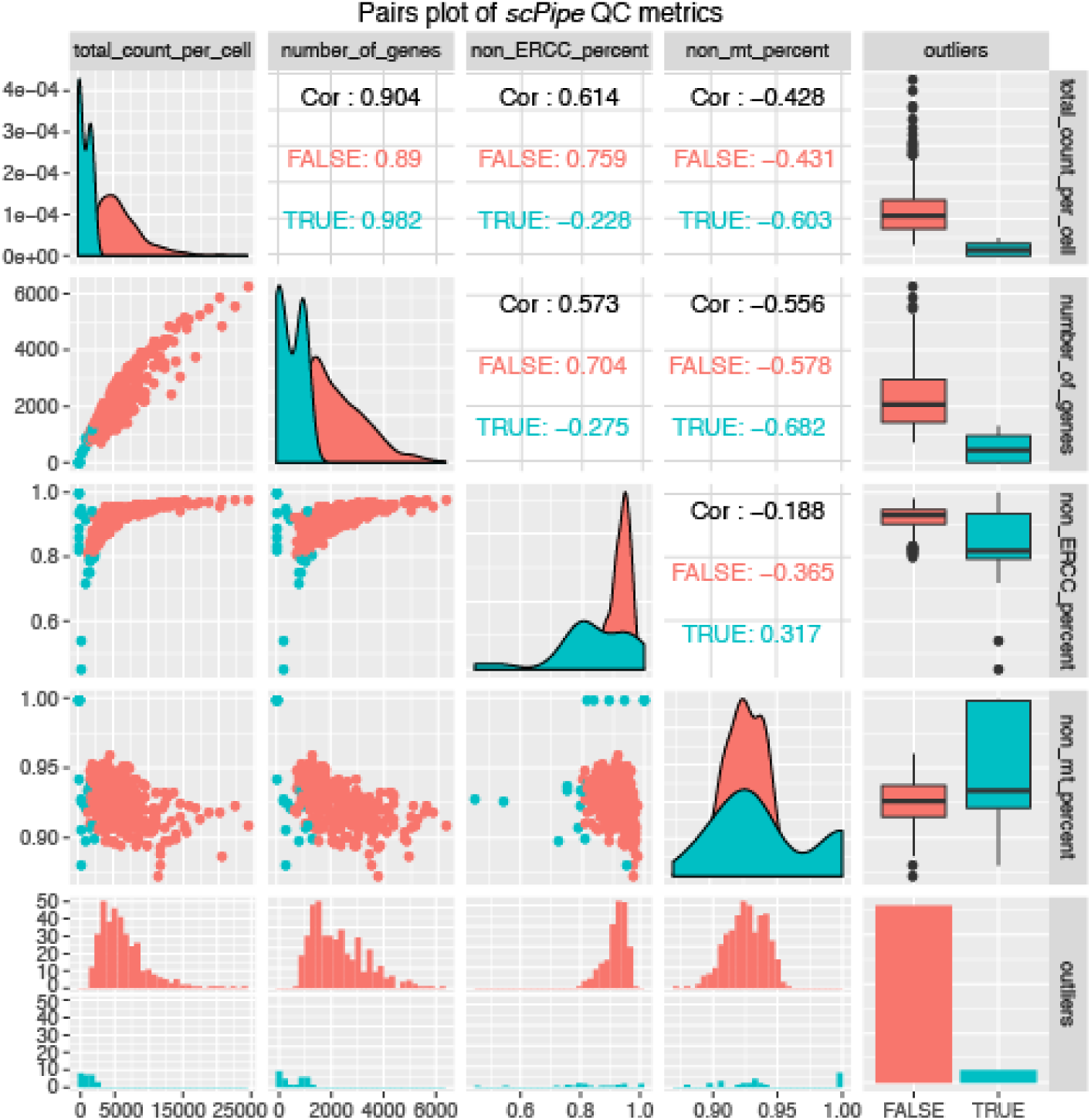
**A pairwise scatter plot of the quality control metrics collected by *scPipe* for the mouse blood cell dataset.** Sample-specific metrics include the total read count, the number of genes detected and the proportion of non-ERCC or non-mitocondrial reads. The good quality and outlier samples detected by *scPipe*’s automatic outlier detection method are indicated in each panel by a different colour.

*scPipe* implements a multivariate outlier detection method for discovering low quality cells and possible doublets to remove from further analysis. The method uses up to 5 metrics (log-transformed number of molecules detected, the number of genes detected, the percentage of reads mapping to ribosomal, or mitochondrial genes and ERCC recovery (where available)) as input (Figure 3). First the gaussian mixture model (GMM) is constructed using these QC metrics to capture the heterogeneity in the dataset. Then for each mixture component, a robustified Mahalanobis Distance is calculated for each cell with outliers automatically detected based on this distance using the *mclust* package [21].

Outlier detection can then be reconciled with visual inspection of the QC metrics through the afore-mentioned QC plotting options to fine tune the sample-specific filtering thresholds chosen. Depending on the data, the only argument that needs to be specified is the maximum component in GMM. In most cases the component is set to 1 for good quality data with only a small proportion of poor quality cells. For data generated by droplet-based protocols where larger proportions of poor quality cells may be expected, the component can be set to 2 to modelling two component which corresponds to the good and poor quality cells. Apart from the number of components, the user can also adjust the confidence interval for choosing outliers. In practice the outlier detection algorithm is fairly robust with regard to this argument. An HTML report generated using *rmarkdown* [14] *that includes all run parameters used, QC statistics and various types of dimension reduction plots of both the gene expression data and QC metrics can be generated using the `create_report` function (Figure 4). As scPipe* generates consistent QC measures across different protocols and experiments, these QC metrics can be easily combined (Figures 2C,2D) to facilitate comparisons between multiple datasets.

**Fig 4.**
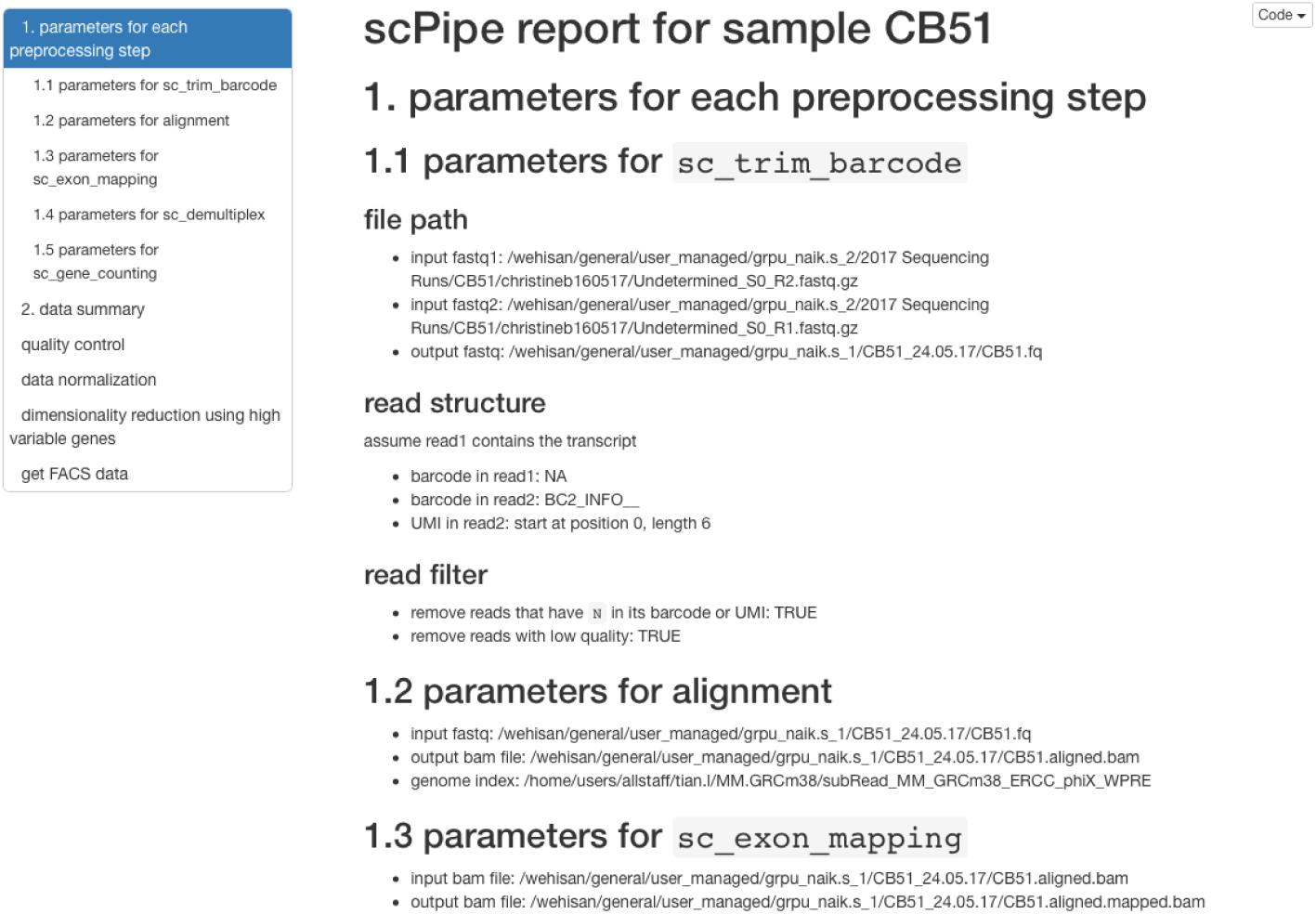
**Screenshot of an HTML report created by *scPipe* for the mouse blood cell dataset.** This report organises the output of *scPipe*, including run parameters and QC metrics and also generates basic dimension reduction plots of the data. Such reports provide a convenient format for communicating basic QC information to collaborators to help them evaluate the overall quality of an experiment.

After quality control, the data can be passed to downstream packages. Use of the *SingleCellExperiment* ensures the output of *scPipe* is fully interoperable with a range of other Bioconductor packages. For instance, packages such as *scran, scater* and *BASiCS* [22] can be used for normalization, *zinbwave* [23] for dealing with zero inflation in the data and removing unwanted variation and SC3 [24] for clustering.

#### Using scPipe

A vignette accompanying the package provides further details on implementation and an example use case on the CEL-seq2 mouse blood dataset. This example dataset contains 100 million reads and takes under 2 hours to process on a standard Linux server. A second example of running *scPipe* on the Chromium 10X cell line data is provided as Supplementary Information (http://bioinf.wehi.edu.au/scPipe/). Processing this dataset which has 400 million reads from around 1,000 cells takes about 10 hours on a standard Linux server. We also provide examples of using *scPipe* with an alternate aligner (STAR [25]). Code for each of these analyses is available from http://bioinf.wehi.edu.au/scPipe/.

#### Comparing *scPipe* and *CellRanger*

In order to highlight the differences between *scPipe* and other tools, we performed a benchmark experiment that included 3 human lung adenocarcinoma cell lines (see Materials and Methods) and processed the raw data using both *scPipe* and *CellRanger* (Figure 5A). The dataset contains about 1,000 cells. *CellRanger* returns 1,027 cells and *scPipe* 981 cells after QC (Figure 5B). The t-SNE plot [26] generated from the *CellRanger* output shows that the 48 cells that appear in the *CellRanger* results but not in *scPipe* tend to cluster together (Figure 5C). Inspection of their QC metrics (Figure 5D) shows that these cells have higher proportions of mitochondrial gene counts, suggesting they may be dead cells that should be excluded from downstream analysis. Since *CellRanger* only uses the UMI counts per cell as a QC cutoff, the results generated by *CellRanger* may contains dead cells, requireing a further round of QC. The *scPipe* analysis on the other hand uses multiple QC metrics by default (Figure 3) to achieve a robust measure of cell quality to ensure low quality cells are discarded. This comparison shows the benefit of *scPipe*’s built-in QC step.

**Fig 5.**
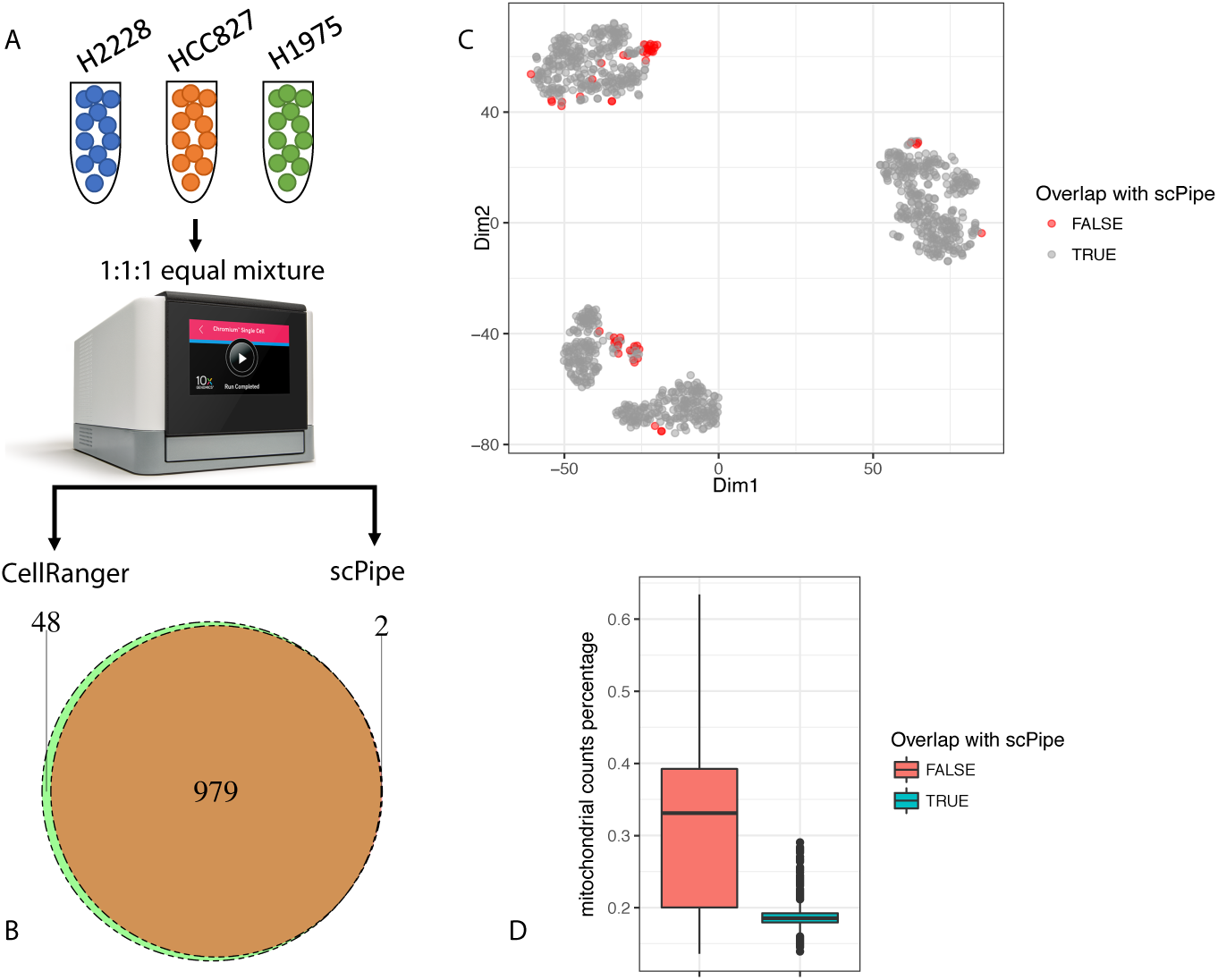
**Comparing *scPipe* and *CellRanger*.** (**A**) The workflow for the comparison. An equal mixture of cells from three cell lines are sequenced using the Chromium 10X platform (see Materials and Methods). Data were processed by *scPipe* and *CellRanger*. (**B**) The pie chart shows the overlap of cell barcode detected by *scPipe* and *CellRanger*. (**C**) The t-SNE plot generated using *CellRanger* output. Cell barcodes that only exist in *CellRanger* are highlighted. **(D)** Box plots showing the percentage of mitochondrial gene counts in cells that overlap with *scPipe* or only exist in the *CellRanger* results.

## Conclusion

With the growing popularity of scRNA-seq technology, many tools have been developed for normalization, dimensionality reduction and clustering. There are relatively few packages designed to handle the raw data obtained from the various 3’ end sequencing protocols with their associated UMIs and cell-specific barcodes from beginning to end and collect detailed quality control informations. The *scPipe* package bridges this gap between the raw FASTQ files with mixed barcode types and transcript sequences and the gene count matrix that is the entry point for all downstream analyses. *scPipe* outputs numerous QC metrics obtained at each preprocessing step and displays these results in an HTML report to assist end users in QC evaluations. Future improvements that are planned for *scPipe* include support for new scRNA-seq protocols as they emerge and parallelization of the various preprocessing steps to enable scalability to larger datasets. We also plan to generate a more comprehensive scRNA-seq benchmark dataset to ensure the default UMI correction and quality control methods used in *scPipe* are optimal.

## Acknowledgments

The authors are grateful to Saskia Freytag, Carolyn de Graaf, Aaron Lun, Rowland Mosbergen, Othmar Korn and Valerie Obenchain for their advice and feedback on the *scPipe* software and Roberto Bonelli for designing the *scPipe* logo.

